# Understanding host response to infectious salmon anaemia virus in an Atlantic salmon cell line using single-cell RNA sequencing

**DOI:** 10.1101/2022.01.04.474990

**Authors:** Ophélie Gervais, Remi Gratacap, Athina Papadopoulou, Ross D. Houston, Musa A. Hassan, Diego Robledo

## Abstract

**Background:** Infectious Salmon Anaemia Virus (ISAV) is an Orthomixovirus that currently represents a large problem for salmonid aquaculture worldwide. Prevention and treatment methods are only partially effective. Genetic selection and genome engineering strategies have potential to develop ISAV resistant salmon stocks. However, this requires a detailed understanding of the genomic regulation of ISAV pathogenesis. Here, we used single cell RNA sequencing on a salmonid cell line to provide a high dimensional insight into the transcriptional landscape that underpin host-virus interactions during ISAV infection at the single cell level.

**Results:** Salmon head kidney 1 (SHK-1) cells were single-cell RNA sequenced before challenge, and at 24h, 48h, and 96h post-ISAV challenge. The results revealed marked changes in the host transcriptome at 48h and 96h post-infection, even in uninfected cells, potentially suggesting paracrine signalling. This paracrine activation of uninfected cells seemed to be unspecific, involving pathways such as mRNA sensing, ubiquitination or proteasome, and also the up-regulation of the mitochondrial ribosome genes. At 24h post infection, cells showed expression signatures consistent with viral entry, with up-regulation of genes such as PI3K, FAK or JNK. At 48h and 96h, infected cells showed a clear anti-viral response, characterised by the expression of IFNA2 or IRF2.

**Conclusions:** This study has increased our understanding of the cellular response of Atlantic salmon during ISAV infection, and revealed potential host-virus interactions at the cellular level. The results highlight the value of single-cell sequencing to characterise cell culture models of viral infection, and the results can be exploited in future functional studies to increase the resistance of Atlantic salmon to ISAV.

## Introduction

Aquaculture is currently the fastest growing food industry worldwide (FAO 2020), and aquaculture products are a fundamental component of healthy and sustainable human diets. Atlantic salmon is the most important aquaculture fish species by value (FAO 2020), and is a high-tech industry underpinned by large research and development programmes. However, infectious diseases remain a major issue for salmon production, such as infectious salmon anaemia (ISA) caused by virulent strains of the ISA virus (ISAV). ISAV is an Orthomyxovirus, and is therefore from the same viral family as Influenza viruses. It is a segmented negative stranded RNA virus, and its genome is formed by 8 RNA segments coding for at least 10 proteins (Rimsad and Markussen, 2019). The disease is characterised by severe anaemia, commonly accompanied by haemorrhage and necrosis of various organs (Aamelfot et al, 2014). ISA disease is listed as notifiable by the World Animal Health Organization, and can cause up to 90% mortality in sea pens. Outbreaks have previously decimated entire national aquaculture industries, as exemplified by the epidemic in Chile (2007-2009, Aamelfot et al, 2014; Godoy et al, 2008). ISAV has been a notifiable pathogen for over 30 years, meaning that upon detection whole stocks have to be culled and the farm is quarantined for a period of time. This culling has major negative impacts on local salmon farms, but also on international trade, exemplified by recent bans on import of genetic material from Norway to the UK.

Preventative and control measures for ISAV include the aforementioned biosecurity measures, and several commercial vaccines also exist. However, these are insufficient to fully control the disease, and improvement of host resistance by selective breeding is an important target. Encouragingly, there is evidence for moderate heritability of resistance to ISA in several studies of Atlantic salmon, and this is being utilised by breeding companies to improve resistance based on regular experimental challenge testing of broodstock families and genomic selection (Kjoglum et al. 2008; Gjerde et al. 2009; Holborn et al. 2020; Gervais et al. 2021). While survival during such challenges is an important target trait, understanding the host-pathogen interactions is important for several reasons. Firstly, it can assist in the identification of functional genetic variants to improve genomic selection accuracy and its persistency across distant relatives. Secondly, it can assist with design of new vaccination, treatment, or management strategies. Thirdly, it can lead to targets for future genome editing studies to potentially develop fully ISA-resistant salmon strains.

Previous studies in Atlantic salmon have reported a notable up-regulation of the innate immune system in response to ISAV (Jorgensen et al. 2008; Lauscher et al. 2011; LeBlanc et al. 2012; Dettleff et al. 2017; Gervais et al. 2021), albeit there is a large degree of tissue specificity in the responses (Valenzuela-Miranda et al. 2015). This is expected since pathogen infections lead to complex and dynamic interactions with the host and its immune system. *In vitro* models provide simplified systems to study the interactions between pathogens and the host cell machinery, which can help break down the multidimensional responses observed *in vivo.* Multiple ISAV infection in vitro studies have been performed, describing a rapid interferon response (Andresen et al. 2020; Samsing et al. 2020) or the interplay between host and virus over the control of oxidative stress and apoptosis (Olavarria et al. 2015a, b). In vitro studies have also been key to understand the role of the different ISAV proteins and their interaction with cellular mechanisms (Li et al. 2016; Zhang et al. 2017; Thukral et al. 2018; Fredericksen et al. 2019; Toro-Ascuy et al. 2019).

While these approaches have provided important insights into ISAV infections in Atlantic salmon, our understanding of this host-pathogen interaction is still incomplete. One of the main limits of previous studies is the measure of population-level cellular responses with limited resolution. Recent advances in single-cell sequencing allows us to obtain a comprehensive picture of the cellular heterogeneity during infection, and the relationship between viral transcription and host molecular responses. Single cell sequencing has yet to be widely applied in Atlantic salmon, but has been used to improve understanding of cellular transcription in gills (West et al. 2021). Here, to better understand the molecular mechanisms of ISAV infection and the Atlantic salmon response to the virus, we have performed an *in vitro* ISAV infection in the salmon head kidney 1 cell line (SHK-1), which has been widely used to study cellular responses to infection. The results reveal the heterogeneity of the infection process and provide a better understanding of the molecular interactions between ISAV and Atlantic salmon.

## Methods

### Cell line

Atlantic salmon head kidney 1 (SHK-1) cells, an immortalised macrophage-like cell line from Atlantic salmon (*Salmo salar*), was obtained from the European Collection of Authenticated Cell Cultures (ECACC-97111106). All cells were grown as a monolayer in L15 complete media (L15*), L15 (Sigma-Aldrich, St. Louis, USA) supplemented with 5% heat-inactivated foetal bovine serum (FBS) (Gibco, Waltham, USA), 40 μM β-mercaptoethanol (Gibco), 100 units/mL penicillin and 100 μg/mL streptomycin (Gibco). Cells were cultured in an incubator at 22 ± 1 °C without CO2. SHK-1 was split 1:2 at 80% confluency with 1/3 conditioned media.

### Viral stock

ISAV stock was obtained from Marine Research (Aberdeen, isolate V4782, derived from UK 2008/2009 outbreak). ISAV was passaged once on 1 x T175 SHK-1 cells (20 mL L15* but with reduced serum (2%) and the supernatant collected after 50% cell death, centrifuged, sterilised using a 0.45 μm filter) and stored in 1 mL aliquots at −80C until use. The viral dose was estimated by flow cytometry. To do this, SHK-1 cells were infected with different doses of the virus for 48 hr in L15* with reduced serum (2%) at 17ºC, followed by antibody staining (ADL anti-ISAv, AquaMab-P10 and anti-mouse-GFP, Invitrogen A21202) and fluorescence quantification (BD Fortessa X20).

### ISAV disease challenge

2.5 x 10^5^ SHK-1 cells were seeded in 6-well plates in L15 + 10% FCS and antibiotics (penicillin and streptomycin) 24h prior to the challenge experiment. The cells were infected with 200 μL of ISAV in L15* with reduced serum (2%) and incubated at 17ºC for 24h, 48h and 96h. Cells (three infected timepoints and control) were collected by trypsin treatment, washed in phosphate buffered saline (PBS, Invitrogen) and suspended to 106 cells/mL in PBS + 0.05% BSA (Bio-Rad cell counter) (Figure 1).

**Figure 1.**
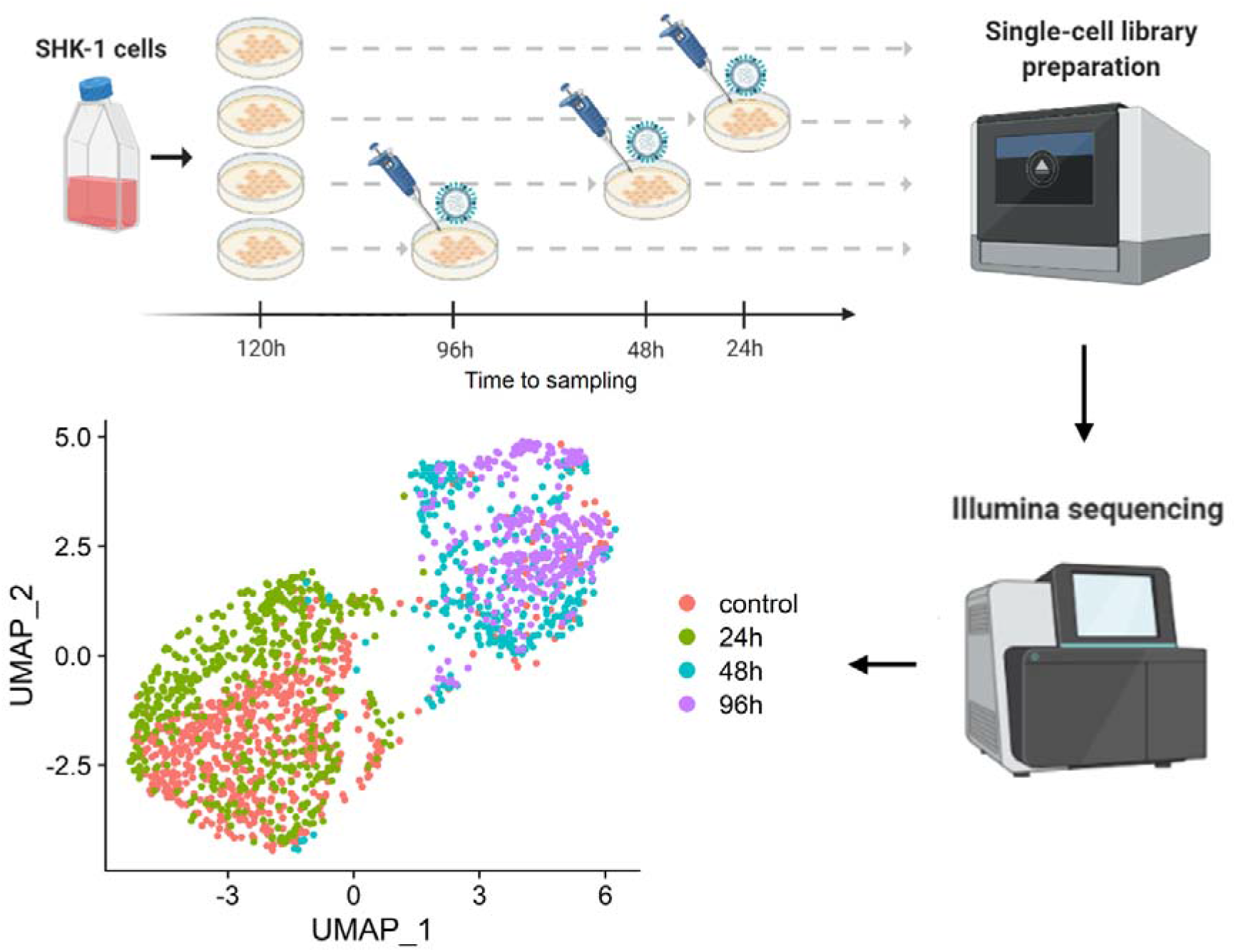
Experimental design and cell clustering.

### Single-cell RNA sequencing library preparation and sequencing

Cells were counted and checked for viability on a Cell Counting Slide (Bio-Rad Laboratories Ltd) and then appropriately diluted according to 10x Guidelines. Each individual sample was loaded separately into the 10x Chromium machine and 10x single cell 3’ GEM kit v3 was used to generate the libraries following the manufacturer’s protocol. The quality of the resulting libraries was assessed in a Bioanalyzer (Agilent) and sequenced in two lanes of a NovaSeq SP flowcell with a cycle setup of 28/8/91 at Edinburgh Genomics.

### Single-cell RNA sequencing analysis

The raw single-cell RNA-seq samples [0h (pre-infection), 24h, 48h and 96h post-infection] were demultiplexed and mapped to a combined reference transcriptome of Atlantic salmon (Ensembl, GCA_000233375.4) and ISAV (NCBI, GCF_000854145.) using Alevin/Salmon v1.4.0 (Srivastava et al. 2019). The resulting raw count matrices were analysed using Seurat v3.1.5 (Stuart at al. 2019) in R v3.6.3 (R Core Team 2017). Each library was loaded individually, discarding genes identified in fewer than 3 cells, and cells with fewer than 200 expressed genes. DoubletFinder V2.0.3 was used to assess the impact of doublets on cell clustering in each library, which was negligible. Thereafter all libraries were merged into a single Seurat object. For each cell, the percentage of mitochondrial and viral transcripts were estimated. After inspection of quality control parameters, cells with fewer than 1,000 or more than 10,000 genes or with percentage of mitochondrial transcripts above 25 were discarded. The infected/uninfected status of all cells was calculated as previously described (Sun et al. 2020) based on the kernel density estimate of the distribution of percentage of viral counts on the log10 scale, which was used to find the first local minima. Cell cycle scores for each cell were estimated using the orthologues of Seurat’s (v3.1.5; Stuart at al. 2019) list of mammalian cell cycle markers. The gene counts of the filtered cells were normalised using SCTransform with the percentage of mitochondrial RNA and cell cycle scores as regression variables. Dimensionality reduction was performed using the Uniform Manifold Approximation and Projection (UMAP) method, and clustered in groups using the ‘FindNeighbors’ and ‘FindClusters’ functions of Seurat (Stuart at al. 2019). Marker genes for each cell group were determined using the Wilcoxon Rank Sum test (logFC > 0.25 and false discovery rate (FDR) corrected p-values < 0.05). Kyoto Encyclopedia of Genes and Genomes (KEGG) enrichment analyses were carried out using KOBAS v3.0.3 (Xie et al. 2011). Briefly, salmon genes were annotated against the KEGG protein database (Kanehisa and Goto, 2000) to determine KEGG Orthology (KO). KEGG enrichment for gene lists was tested by comparison to the whole set of expressed genes (obtained from the Seurat object) using Fisher’s Exact Test. KEGG pathways with ≥ 5 DE genes assigned and showing a Benjamini-Hochberg FDR corrected p-value < 0.05 were considered enriched.

## Results

To assess the intracellular activity of ISAV and study the corresponding host response, single-cell RNA-Seq was performed on SHK cells infected with ISAV 24h, 48h and 96h post infection (Figure 1). Uninfected SHK cells were used as controls. Raw reads were assigned to either Atlantic salmon genes or ISAV genes using a combined host-virus reference transcriptome. A total of 638, 512, 296 and 323 cells passed quality control filters for the control, 24h, 48h and 96h samples respectively. Clustering of the cells in these four libraries based on the host transcriptional response showed a clear separation between control & 24h infected samples and the 48h & 96h infected samples (Figure 1). A small fraction of the cells showed an unexpected behaviour (i.e. clustering together with cells from a different and unexpected timepoint), but this can be attributed to index hopping and cross-contamination with ambient mRNA (Farouni et al. 2020).

### Viral transcription during the infection of Atlantic salmon cells

The percentage of infected cells (estimated following Sun et al. 2020) was 16% at 24h post-infection, and 32-33% at 48 and 96h (Figure 2A). The infected cells formed two different groups, one with a small number of 24h infected cells, and the largest one with 48h and 96h infected cells, where the percentage of viral transcripts was markedly higher, representing more than 20% of the transcriptome of the cells (Figure 2B). Expectedly, the percentage of virus transcriptome, relative to the total transcriptome increased over the course of infection, mostly ranging between 2% and 15% (Figure 2C). This is consistent with previous reports in other orthomyxovirus, although a larger heterogeneity has been described, with viral transcript levels reaching up to 90% of the cell transcriptome (Russel et al. 2018; Sun et al. 2020). Viral transcript levels can vary substantially depending on the cell type (Sun et al. 2020), but also ISAV infection progression is typically slower than that of other orthomyxovirus, thus, later timepoints could show higher ISAV transcript levels.

**Figure 2.**
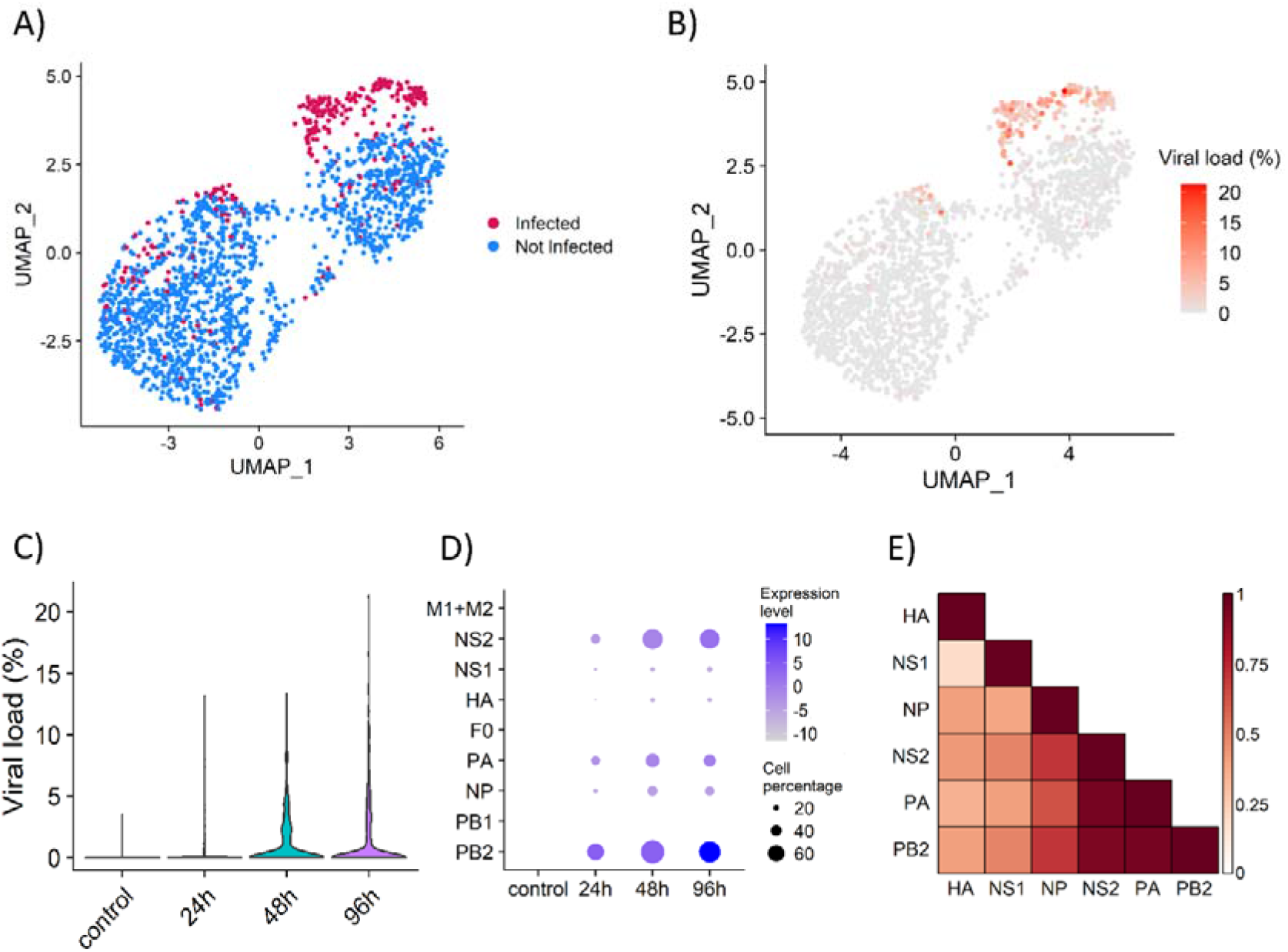
ISAV infection and transcription. A) UMAP dimensionality reduction plot showing the infected and uninfected cells. B) UMAP dimensionality reduction plot showing the viral load of each cell measured as percentage of viral transcripts. C) Viral load in each sample, measured as percentage of viral transcripts. D) Dotplot showing the expression level and the percentage of host cells with transcripts of each viral gene in each sample. E) Heatmap showing the correlation between the expression of the viral genes.

The timing and level of expression of each viral gene also varies along the infection process (Figure 2D); the viral polymerase basic protein 2 (PB2) gene is the most expressed throughout the infection, followed closely by the polymerase acidic protein (PA) and the non-structural proteins 1 (NS1) and 2 (NS2, also termed nuclear exporting protein or NEP). NS1 and NS2 were measured together to avoid the 3’ bias of 10x single-cell RNA-Seq technology; NS2 is a splicing product of NS1 (Ramly et al. 2013) and is expressed at lower levels according to previous reports in ISAV (Ramly et al. 2014) and Influenza A (Robb et al. 2010). The expression of these three genes showed a high positive correlation (Figure 2E), suggesting their co-expression early during the infection. Expression of the hemagglutinin (HA) is found in a small fraction of cells, and it shows the lowest correlation with the other (expressed) viral genes (Figure 2E). Expression of the bicistronic mRNA encoding the matrix protein (M1) and a nuclear export protein (M2, also termed S8ORF2), of the polymerase basic 1 (PB1) and the viral membrane fusion protein (F0) was not detected in our dataset (Figure 2D), which suggests they are either expressed late during the infection or expressed at low levels. In Influenza A all the viral mRNAs show similar transcription patterns (Phan et al. 2021), however as mentioned above, ISAV infection typically progresses at a slower rate so it is possible that the viral mRNA dynamics are slightly different and easier to detect.

### Host response to ISAV

The cells were clustered in 8 different groups according to the host gene expression (Figure 3A). Clusters 1 to 4 represent the control and 24h samples; most control cells are part of clusters 1, 2 and 3, which also have many 24h cells, while cluster 4 is composed mostly of 24h cells with an important number of infected cells. Clusters 6 to 8 represent the 48h and 96h samples; cluster 6 is formed by 48h and 96h uninfected cells, while cluster 7 and 8 are 48h and 96h infected cells, respectively. The clear separation of control-24h and 48h-96h uninfected cells suggests paracrine regulation by infected cells. Paracrine signalling has been described before in response to Influenza A (Ramos et al. 2019) and other viruses (Patil et al. 2015; Voigt et al. 2016), eliciting an antiviral interferon-mediated response in uninfected cells. Finally, cluster 5 is the most difficult to interpret since it is formed by a low number of cells of each sample apparently not connected to the infection process (Figure 3A).

**Figure 3.**
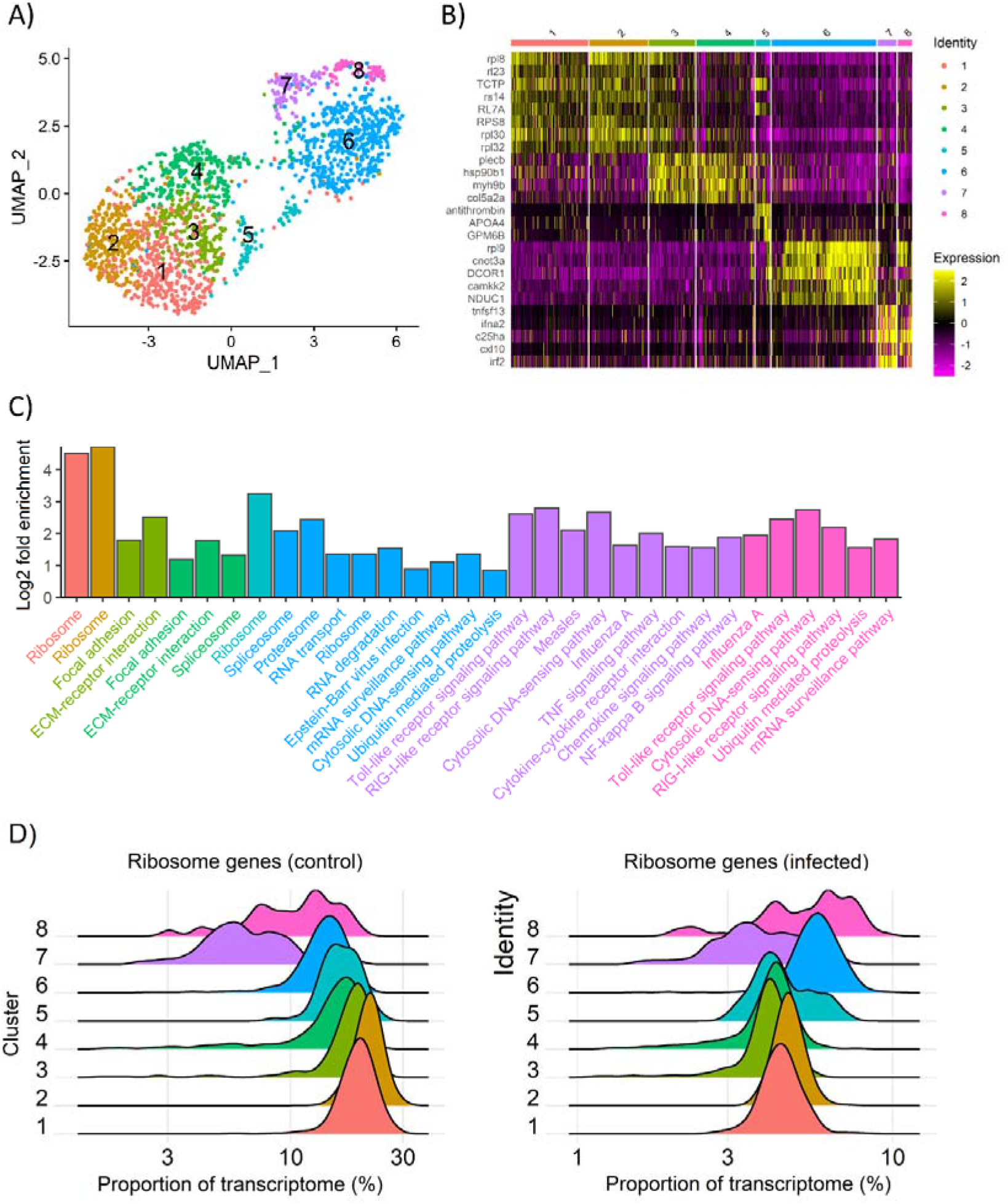
Response to ISAV in the Atlantic salmon SHK-1 cell line. A) UMAP dimensionality reduction plot showing the clustering of the cells according to the host cell transcriptome; B) Heatmap showing the expression pattern of top marker genes of the UMAP cluster; C) Barplot with the fold enrichment of selected KEGG pathways in each cell cluster; D) Ridgeplot showing the proportion of the transcriptome that represent the genes assigned to the enriched KEGG pathway “Ribosome” in cluster 1 and cluster 5.

Each of these clusters is characterised by specific marker genes (Figure 3B, Supplementary File). Clusters 1 and 2 are characterised by the high expression of certain ribosomal genes, which also show a high expression in half of the cells of group 3. This suggests manipulation of the ribosomal machinery by ISAV, which can be connected to the hijacking of host translational machinery by the virus, a common strategy used by virus to repress the cellular mRNA translation and allow the preferential translation of viral mRNAs (Levene and Gaglia, 2018). On the other hand, a group of immune genes, including interferon alpha 2 (*Ifna2*) or interferon regulatory factor 2 (*Irf2*), are highly expressed in infected cells at 48h (cluster 7), and moderately in infected cells at 96h (cluster 8), suggesting either viral repression or a shift in the immune response later during the infection. Further, this cluster is not highly expressed in non-infected cells at 48h and 96h (cluster 6), suggesting that paracrine signalling by infected cells does not promote the activation of an interferon response in bystander cells.

KEGG pathway analysis (Figure 3C) confirmed the enrichment in ribosomal genes in the uninfected cells clusters from early timepoints (1 and 2). Clusters 3 and 4 (mostly 24h cells, especially cluster 4) are enriched for “focal adhesion” and “ECM-receptor interactions”, likely a consequence of the interaction of ISAV with the extracellular matrix and membrane receptors, and successive internalization. The genes in these pathways include phosphoinositide 3-kinase (PI3K), focal adhesion kinase (FAK) or c-Jun terminal kinase (JNK), important for Influenza A entry into the cytoplasm (Ayllon et al. 2012; Elbanesh et al. 2014; Zhang et al. 2018). This is consistent with the similarity in the entry mechanisms of Influenza A and ISAV (Eliassen et al. 2000; Kibenge and Kibenge, 2016) and suggests that molecular interactions during viral entry are also conserved to some extent. These early genes are good candidates for functional studies aimed at disrupting ISAV entry into salmon cells.

Uninfected cells at 48h and 96h (cluster 6) showed a non-specific state of alert, with terms such as “proteasome”, “RNA degradation”, “mRNA surveillance pathway” or “cytosolic DNA-sensing pathway”. For instance, this cluster shows expression of the seven lsm genes that compose the Lsm1-7 ring, involved in the degradation of messenger RNA in the cytoplasm (REF). Many proteasome-specific genes are also expressed in this cluster, including two of the subunits of the immunoproteasome, involved in the degradation of intracellular proteins, including those of viral origin (REF). Connected to the proteasome, this cluster also expresses all the genes of the ubiquitination machinery: E1 ubiquitin-activating enzymes, E2 ubiquitin-conjugating enzymes and E3 ligases of all the types (HERC3, U-box and RING-finger and their adaptor proteins). Spliceosome genes are also enriched, suggestive of active post-transcriptional and post-translational mechanisms in this cluster.

Uninfected cells at 48h and 96h (cluster 6) also showed enrichment in ribosomal genes, as uninfected control and 24h cells (cluster 1 and 2), which was unexpected based on the expression pattern of ribosomal genes in the heatmap (Figure 3B). However, we found that the underlying specific ribosomal genes are different, with only two genes in common between the two groups of cells (out of 55 in each list). These two sets of ribosomal genes show contrasting patterns of expression (Figure 3D), with the “control” ribosomal genes showing less expression in infected cells, while the “infected” ribosomal genes show increased expression at 48h and 96h. A closer look at the gene lists revealed that most of the “infected” ribosomal genes code for proteins of the mitochondrial ribosome (Supplementary file X). Mitochondria activate anti-viral immune responses through mitochondrial antiviral signalling (MAVS; Refolo et al. 2020) and can also initiate apoptosis (Lei et al. 2009). Influenza A NS1 protein has been observed in the mitochondria (Tsai et al. 2017), and is able to alter mitochondria morphodynamics (Pila-Castellanos et al. 2021). Our results suggests that paracrine regulation upon ISAV infection activates the mitochondria as a defence mechanism, and the fact that these mitochondria ribosomal genes are also overexpressed at 96h in infected cells (Figure 3D, cluster 8) suggests that they play a role in the immune response against ISAV.

Finally, clusters 7 and 8 (infected 48h and 96h cells) show up-regulation of key immune response genes, such as interferon alpha 2 and 3 (ifna2, ifna3), interferon regulatory factor 2 (irf2) or C-X-C motif chemokine 10 (cxl10). These genes are slightly more up-regulated in the 48h cells group (Figure 4), with pathways such as toll-like receptor signalling, RIG-I receptor signalling, TNF signalling or cytokine-cytokine interaction. However, these genes were not expressed in uninfected cells at 48 and 96h (cluster 6). IFNA2, IFNA3 and CXL10 are secreted proteins that act on other cells to promote an antiviral state and could explain the “activated” state of cluster 6 cells. However, surprisingly they would not induce the up regulation of interferon genes.

**Figure 4.**
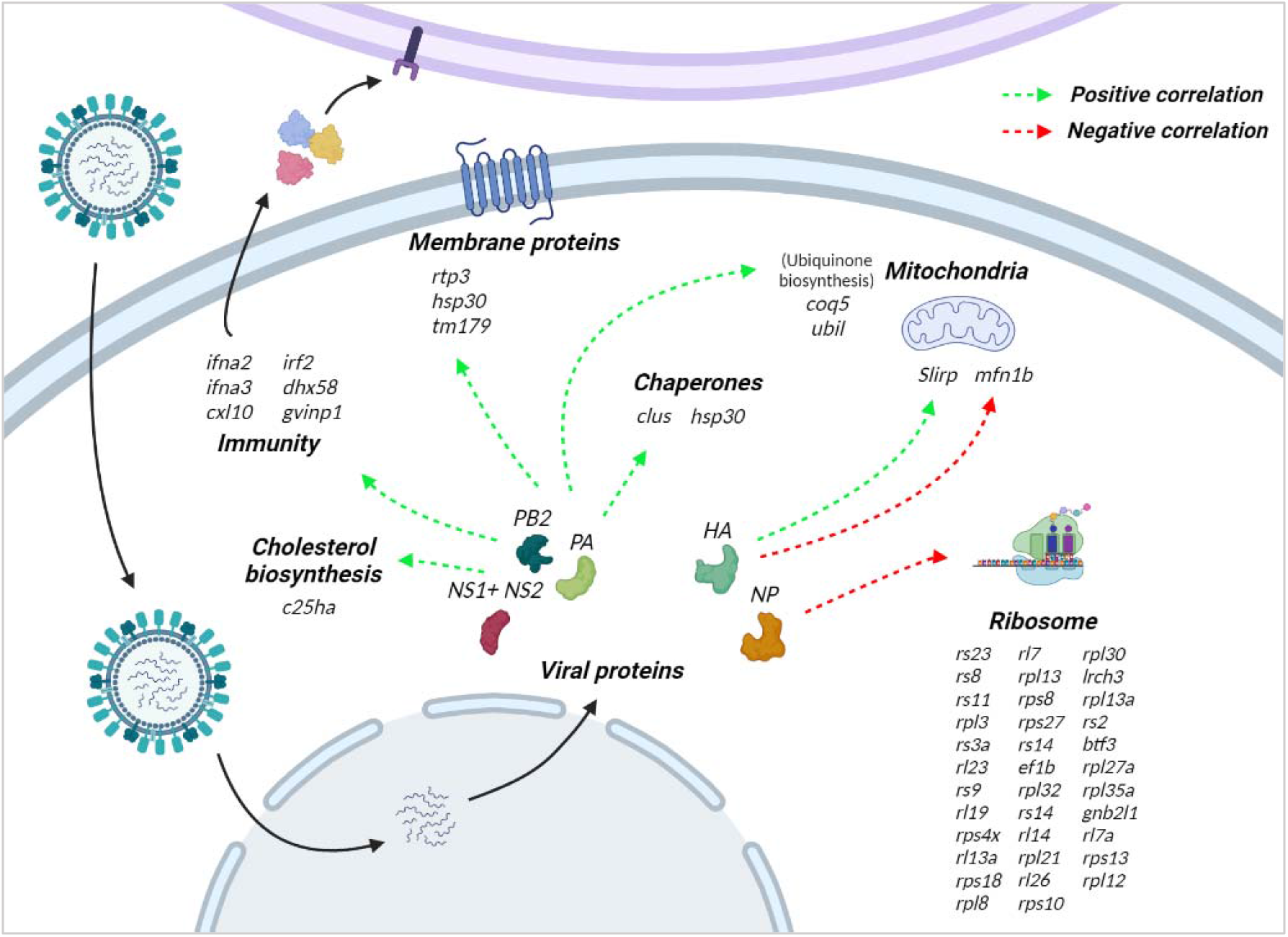
Correlation between viral gene expression and host gene expression.

### Interactions between host and viral transcripts

To evaluate whether specific viral proteins could have a direct impact on the expression of host genes, a correlation analysis between viral and host genes was performed for infected cells (Figure 4, Supplementary file). *PB2*, *PA* and *NS1+NS2* expression patterns showed significant correlation with largely the same set of host genes (r > |0.3|), as expected considering the highly correlated expression of these viral genes. Considering these viral genes are the most expressed, these correlations do not necessarily represent regulation of host genes by viral proteins, they could also represent genes that respond proportionally to viral load. In fact, the host genes exhibiting the highest positive correlation include the immune genes mentioned in the previous section (*ifna2, ifna3, irf2*) and the regulator of antiviral signalling probable ATP-dependant RNA helicase DHX58 (*dhx58*). The highest correlation was observed with cholesterol 25-hydroxylasel A (*c25ha*), an interferon-stimulated gene involved in the regulation of cholesterol biosynthesis, which has broad antiviral activity (Liu et al. 2013). However, these viral genes, especially *NS1+NS2*, also show negative correlations with several transcription factors, such as twist-related protein 2 (*twst2*) or zinc-finger protein SNAI1 (*snai1*).

Consistent with previous results, the expression of over thirty cytoplasmic ribosomal genes is negatively correlated with the expression of the viral nucleoprotein NP (r = −0,3 to −0.4) (Figure 5), suggesting a negative regulation of the host translation machinery by the virus. While NP is a structural protein, it has been shown that it interacts with host factors to facilitate viral replication (Momose et al. 2001; Kawaguchi et al. 2011), however no interactions with ribosomal genes have been previously described. There were also several chaperons showing positive correlation with viral genes, such as clusterin (clu), which has antiapoptotic activity and is directly targeted by the Influenza A virus nucleoprotein (Tripathi et al. 2013). Another protein of interest is sequestosome-1 (sqstm1), a receptor required for selected autophagy that inhibits Seneca valley virus and avian influenza replication (Wen et al. 2021; Liu et al. 2021).

## Discussion

In this study single-cell RNA sequencing was applied to study ISAV infection in Atlantic salmon SHK-1 cells. This simplified system, in combination with this novel genomics technology, has allowed us to finely characterise the transcriptomic changes that occur during ISAV infection both in the virus and the host, and suggest potential interactions. While cell line models are often used to study hostpathogen interaction, this study highlights that even within an immortalised cell line there is major heterogeneity in both infection status and cellular immune response. Our study complements and expands knowledge gained in previous *in vivo* and *in vitro* studies of ISAV infections in Atlantic salmon.

Our results align with previous studies that have shown that ISAV triggers an early immune response characterised by the activation of interferon genes, both *in vitro* and *in vivo* (Jorgensen et al. 2007; Svingerud et al. 2013; Valenzuela-Miranda et al. 2015). An interesting finding is that the interferon response was only triggered in infected but not in bystander cells, which is quite different to what has been reported during Influenza infections, where the paracrine signalling is important at 12h post infection, increasing the expression of interferon stimulated genes in bystander cells (Ramos et al. 2019). There are different potential explanations for this finding, for instance the virus may be able to inhibit the interferon-related paracrine signalling quite early during the infection, but it is also plausible that paracrine signalling functions differently in fish and mammals. Further investigations are necessary to understand fish paracrine signalling, including studies with other species and viruses.

The ribosomal machinery is an important part of viral replication since viral genomes do not usually harbour mRNA translation genes, and therefore they need to recruit host proteins including ribosomal proteins (RPs) to complete their replication cycle. Our result shave shown that the viral protein NP may interact or at least drive the expression of specific mitochondrial RPs. While no interactions between NP and ribosomal proteins have been described, it is well known that viruses interact with ribosomal proteins as part of their infection strategy (Li 2019; Dong et al. 2020). However, recent studies have also highlighted that some ribosomal proteins can have an antiviral function by either interacting with viral proteins to inhibit transcription/translation or by activating antiviral defence signalling pathways (Li 2019). Non-infected cells in 48h and 96h infected samples showed a clear up-regulation of mitochondrial RPs, which could suggest that they form an important part of an antiviral state pathway in bystander cells, potentially repressed by the viral protein NP in infected cells considering the negative correlation.

The proteasome and ubiquitination also seem to be an important component of the antiviral state in bystander cells. We have also found up-regulation of genes related to ubiquitination in resistant fish in response to ISAV (manuscript in preparation). Moreover, the protein encoded by ISAV segment 8, which is known to interfere with interferon signalling, can be conjugated to ubiquitin and the ubiquitin-like interferon simulated gene 15 (ISG15) is Atlantic salmon cells (Olsen et al. 2016). Several viruses can hijack the host ubiquitination machinery, and in particular Influenza uses host ubiquitination as part of its cell entry and replication strategy (Rudnicka and Yamauchi, 2016; Huang et al. 2018; Gu and Fada, 2020).

Our study has highlighted significant correlations between viral and host genes, including several chaperones. Chaperones are used by viruses to facilitate the folding and assembly of the viral proteins (Aviner and Frydman, 2022). There are also host genes associated to autophagy process that are correlated with viral genes. Previous studies have reported that mammalian cells can use autophagy to restrict the replication of avian Influenza (Liu et al. 2021), but it has also been reported that it can promote the replication of influenza A (Wang et al. 2019; Zhang et al. 2021). In any case, autophagy-related genes and process may represent novel targets for the development of anti-ISAV therapeutics.

## Conclusion

Early ISAV infection of Atlantic salmon SHK cells is characterised by high expression of the PB2, NS1+NS2 and PA viral genes, which show highly correlated expression patterns. After 24h of infection infected SHK cells showed similar transcriptomic profiles to control cells, however genes and pathways connected to viral entry were identified, which could be good targets for functional studies to impair viral infection. At 48h and 96h post-infection there was a clear host transcriptomic response to the virus which also affected uninfected cells, therefore suggesting paracrine signalling from infected cells. These paracrine signalling seemed to produce an unspecific alert state in the uninfected cells, involving mRNA surveillance and RNA degradation, ubiquitination and the proteasome, or mitochondrial activation. These pathways were also up-regulated in 48h and 96h infected cells, but they also showed a clear antiviral interferon response, including genes such as *ifna* or *irf2*. Correlation between the expression of viral and host genes suggested potential negative regulation of certain host transcription factors and ribosomal genes by viral proteins. This study increases our understanding of the molecular interactions between ISAV and Atlantic salmon cells, and provides new targets for functional studies aiming to increase the resistance of Atlantic salmon stocks, leading to increased food security and fish welfare.

## Acknowledgements

This work was supported by an RCUK-CONICYT grant (BB/N024044/1) and BBSRC Institute Strategic Funding Grants to the Roslin Institute (BBS/E/D/20002172,BBS/E/D/30002275, and BBS/E/D/10002070). Sequencing was carried out by Edinburgh Genomics, which is partly supported through core grants from NERC (R8/H10/56), MRC (MR/K001744/1), and BBSRC (BB/J004243/1).

## Notes

### Competing Interest Statement

The authors have declared no competing interest.

